# Successful single-session neural self-regulation through neurofeedback varies between features

**DOI:** 10.64898/2026.01.07.698228

**Authors:** E Syrjänen, J Silva, E Astrand

## Abstract

Neurofeedback (NFB) and Brain-Computer Interface (BCI) research seldom present within-session individual learning dynamics. This is even though a large proportion of NFB and BCI users cannot learn the neural self-regulation required to control the feedback. Understanding the time course and learning dynamics between subjects will enable us to design more effective NFB and BCI protocols that promote the learning of neural self-regulation. In this study, we aimed to analyze individual learning trajectories of self-regulation of four different cortical rhythms, in terms of both frequency and spatial selectivity. Twenty healthy subjects performed four sessions of NFB training, each session with feedback reflecting a different cortical rhythm as measured with an electroencephalogram. We specifically tested frontal midline (fm) Theta, occipital Alpha, unilateral centrotemporal sensorimotor rhythms (SMR), and central Beta. We show that all subjects were able to self-regulate at least two of these features, however, with varied specificity in the spatial and frequency domains. Unexpectedly, we show that none of the subjects succeeded in regulating fm Theta. Using a clustering approach, we identified two different learning dynamics among the learners across features: a linear increase/decrease and a non-linear plateau-like trajectory. This is the first NFB study employing an intra-subject cross-over experimental design, enabling the direct comparison of neural self-regulation between multiple features. Our results provide important insights into the “non-learner” problem, showing that it is not a feature-universal personal trait. We further show feature-specific spatial and frequency selectivity of neural self-regulation, providing important considerations for future NFB protocols.

## Introduction

The human brain, a product of millions of years of evolution, exhibits remarkable capabilities, sustaining complex thought, processing vast amounts of information, and controlling the intricate processes that regulate our bodies, all while consuming energy comparable to a few LED lights. The fundamental operation of the brain relies on synchronized neuronal activity, both within local neuronal populations and across distributed brain networks (Buzsaki & Draguhn, 2004; Buzsaki et al., 2013). This synchronized activity, measurable via electroencephalography (EEG) when sufficiently large neuronal populations fire in unison, can be decomposed into distinct frequency bands, each associated with a range of cognitive functions (Basar et al., 1999; Klimesch, 1999). The brain’s capacity for flexible information processing hinges on a delicate balance between synchronized and desynchronized neuronal activity (Engel et al., 2001; Ros et al., 2014). A general principle is that higher frequencies, such as the Gamma band (30-100 Hz), are associated with bottom-up encoding of sensory information and perceptual binding (Fries, 2009; Singer, 1999). In contrast, lower frequencies, like frontal Theta (4-8 Hz), are implicated in top-down control processes, coordinating and modulating the encoding of information (Lisman & Jensen, 2013; Womelsdorf et al., 2006). Frontal Theta, in particular, has been linked to a variety of cognitive control functions, including working memory, error monitoring, and conflict resolution (Cavanagh & Frank, 2014; Cohen, 2014). Similarly, other frequency bands, such as parieto-occipital Alpha (8-12 Hz), has been related to attentional processes and sensory gating (Jensen & Mazaheri, 2010; Klimesch, 2012), and sensorimotor rhythms (SMR; 12-15 Hz), have been associated with motor preparation and inhibition (Pfurtscheller & Lopes da Silva, 1999).

In the 1950s, Joe Kamiya’s pioneering introspection experiments revealed that individuals could consciously alter their brain states, specifically their Alpha activity, in anticipation of reporting their subjective experience (Kamiya, 2011). Subsequent experiments demonstrated that participants could learn to voluntarily control their Alpha activity through feedback (Kamiya, 1968). In the 1960s, Sterman and colleagues showed that cats trained to increase SMR activity not only exhibited reduced seizure susceptibility but also retained this enhanced activity even one month after training had ceased (Sterman et al., 1970; Sterman & Wyrwicka, 1967). These seminal discoveries established the foundation of neurofeedback (NFB) training, demonstrating the potential for individuals to gain volitional, long-lasting control over their brain activity through practice. The causal relationship between NFB training and behavior was first reported in humans a couple of years later (Kaplan, 1975; Sterman & Friar, 1972; Sterman et al., 1974).

The proposed mechanism underlying NFB training is operant conditioning, or reinforcement learning (Koralek et al., 2012; Sitaram et al., 2017). When participants receive positive feedback contingent upon the modulation of a specific brain activity pattern, the positively reinforced neural activity is more likely to be repeated. As participants explore different mental strategies, the feedback they receive fluctuates, guiding them towards the targeted brain state. Positive feedback is associated with the release of dopamine, a neurotransmitter that promotes synaptic plasticity, thus consolidating a “memory” of the brain activity that elicited the reward (Ros et al., 2014; Schultz, 2002). Through this iterative process, participants gradually learn to volitionally regulate their brain activity (Gruzelier, 2014).

The clinical potential of NFB training becomes apparent when considering that many non-neurodegenerative neuropsychiatric disorders are characterized by altered patterns of synchronized and desynchronized brain activity compared to healthy populations. Theoretically, NFB could serve as a non-invasive method for “retuning” brain activity towards a more normative state. Several meta-analyses and review articles suggest that NFB training can lead to positive outcomes in conditions such as stroke (Renton et al., 2017), attention-deficit/hyperactivity disorder (ADHD) (Arns et al., 2009), post-traumatic stress disorder (PTSD) (Panisch & Hai, 2020), and obsessive-compulsive disorder (OCD) (Ferreira et al., 2019). However, these reviews also highlight significant methodological challenges in the field, including a lack of specificity and difficulty in differentiating treatment effects from placebo, regression to the mean, or spontaneous remission (Thibault & Raz, 2017). Even in well-controlled, double-blind randomized controlled trials (RCTs), providing sham feedback to control groups can negatively impact motivation, as participants may not perceive the feedback as rewarding (Ramot & Martin, 2022). Furthermore, translating promising research findings into clinically viable therapies has proven difficult, partly due to the heterogeneity of symptoms and co-morbidity with other psychiatric disorders (Hammond, 2010; Micoulaud-Franchi et al., 2014). Despite these challenges, the potential benefits of NFB, including its safety (Hammond, 2010) and the possibility of long-lasting therapeutic effects, offer a compelling alternative or adjunct to traditional pharmacological treatments, which can be associated with adverse side effects and often require lifelong adherence.

A persistent challenge in NFB and brain-computer interface BCI research is the phenomenon of “illiteracy” where a significant proportion of individuals (estimated at 16-57%) fail to achieve meaningful control over their brain activity (Alkoby et al., 2018; Sannelli et al., 2019; Vidaurre & Blankertz, 2010). While demographic and personality factors have not consistently predicted NFB performance (Weber et al., 2020), it is increasingly recognized that individual differences in brain activity, learning styles, and task strategies play a critical role (Kober et al., 2013). The diversity of training protocols is a crucial factor contributing to the variability in NFB and brain-computer interface (BCI) performance (Sadtler et al., 2014). In a study with Rhesus macaques, Sadtler et al. (2014) demonstrated that individuals have inherent “neural degrees of freedom,” suggesting that certain brain activity patterns are more readily modifiable than others. This raises a fundamental unresolved question: whether the widely reported “non-learner” phenomenon reflects a stable, individual trait or whether it depends on the specific neural feature being targeted. Critically, most prior studies investigate a single neurofeedback feature per cohort, making it difficult to disentangle inter-individual variability from feature-specific effects. As a result, it remains unclear whether individuals who fail to regulate one neural signal would also fail under different neurofeedback targets, or whether learnability is context-dependent.

To address this gap, the present study employs an intra-subject design in which each participant performs neurofeedback across four distinct EEG features (frontal midline Theta, occipital Alpha, centrotemporal SMR, and central Beta) under controlled conditions. This approach enables a direct comparison of neural self-regulation across features within the same individuals, allowing us to test whether variability in performance reflects stable individual differences or feature-specific constraints. We hypothesized that neurofeedback performance would vary substantially within individuals across features, rather than reflecting a uniform ability or inability to learn.

## Materials and Methods

### Participants

Twenty healthy adults (15 males and 5 females) aged between 20 and 39 (M = 27.60, SD = 5.57) participated in one session of computerized cognitive tasks and four sessions of real-time EEG-based NFB training. The participants trained on one different NFB feature in each session. Specifically, sessions randomly alternated between frontal midline Theta (4–7 Hz), occipital Alpha (8–12 Hz), centrotemporal SMR (13–15 Hz), and central Beta (18–25 Hz). Table I expands on each feature configuration. The study adhered to the principles of the Declaration of Helsinki and complied with local rules and regulations, as approved by the Swedish Ethical Review Authority (Dnr. 2021-03121).

**Table I.**
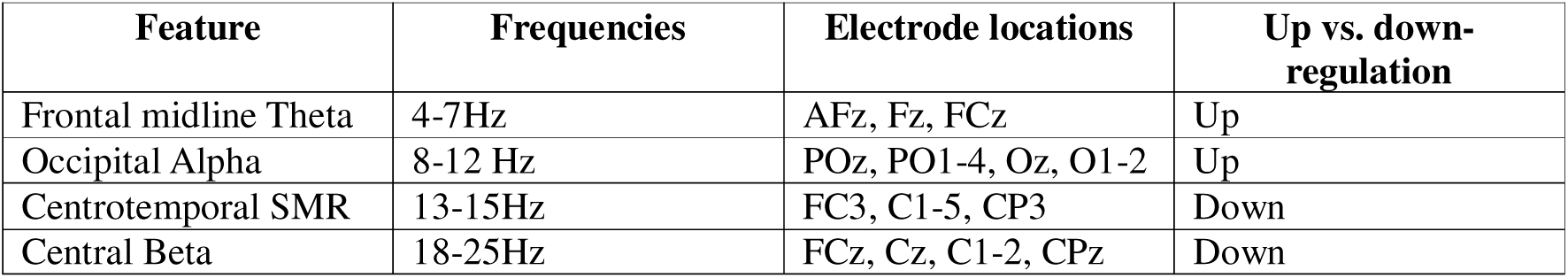
NFB feature specifics.

### Measurements

The real-time NFB system streamed data to a custom-made LabVIEW environment. Data packets of EEG data from LabVIEW were assembled into pre-defined windows of 250 ms that overlapped by 150 ms and were sent to MATLAB for signal processing (using custom-made MATLAB scripts).

### Overall protocol

Participants performed various computerized cognitive tasks in the first session (i.e., working memory, executive functioning, covert visual attention, and multitasking). Considering the scope of this study, the results from the first session are not reported here. In the following four sessions, the participants trained at controlling one individual feature, with each session focusing on a different feature. The features were: up-regulation of frontal midline Theta, up-regulation of occipital Alpha, down-regulation of centrotemporal SMR, and down-regulation of central Beta. The order of features was randomly assigned before the first session. The selection of features and regulation directions was guided by well-established relationships between EEG frequency bands, their associated cortical regions and related cognitive functions, aiming to span a diverse range of cognitive states, as outlined in the Introduction.

Following the completion of each session, participants were verbally debriefed by the experimenters. To gather insight into the cognitive processes employed during the task, participants were probed with a series of open-ended questions, such as: “What were you thinking to try to control the arrow?” and “Did you find any specific strategy that seemed to work?”. We documented the participants’ responses in a logbook. This procedure was conducted for each of the four NFB sessions per participant. Due to operator error, strategy reports for a small number of sessions were not recorded.

### Neurofeedback experiment

During the NFB training, participants were seated in front of a computer screen and instructed to make an arrowhead point as much upwards as possible on the screen (Figure 1). No strategies for controlling the arrowhead were provided to the participants, and they were instructed to control it mentally without using any muscle activity. One session consisted of 64 trials, each lasting 30 seconds. Between each trial, there was a 5-second pause with instructions displayed and a 1.5-second preparation period. Every eight trials, there was a longer 2-minute pause. After 16 minutes of NFB training (halfway through the session), there was a long break for rest and snacks. Each trial began with an instruction, and before the trial started, the instruction was replaced by a grey vertical line that turned white at the start of the trial. See Fig. 1 for an example of the trial structure. The arrowhead’s direction and pointiness reflected the average power magnitude across frequencies and channels for each feature (see Table I. for each feature’s specifics). The direction of the arrowhead would depend on whether power was above or below a threshold. For up-regulated features, the arrowhead pointed up when power was above the threshold. For down-regulated features, the direction of the arrowhead was reversed so that the arrowhead pointed up when power was below the threshold (i.e., the participants always received positive feedback as the arrowhead pointed upwards). The pointiness was determined as the ratio between the current power and the threshold. The threshold was initialized to a constant value of .2 that was used for the first time windows of NFB. During the first eight trials (each 30 seconds long), the threshold was recomputed and adapted every second based on the median of the 95th percentile of the power values. For the remainder of the session, the threshold was fixed to the median of the 95th percentile of the power values from the first eight trials. Using a photodiode attached to the screen, we calculated the delay between the EEG activity (last sample in the 250-ms window) and the display of feedback to range between 70 ms and 80 ms (computed with a method similar to Belinskaya et al., 2020).

**Figure 1.**
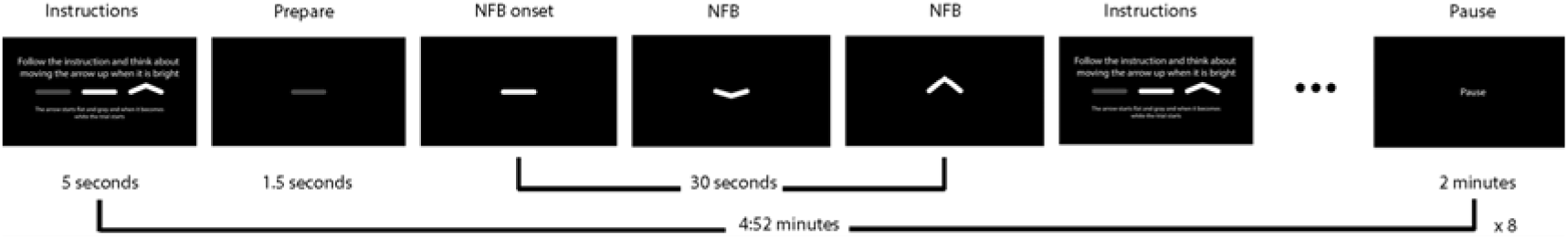
Task instructions and overview of a NFB session. At the beginning of each NFB trial, the instruction was shown for 5 seconds, followed by a horizontal grey line displayed for 1.5 seconds. NFB task onset started as the line became white and lasted for 30 seconds. NFB was continually provided by the orientation and pointiness of the line, updated every 100 ms. Positive feedback (of successful self-regulation) was provided by the arrowhead pointing upwards. Each trial ended by the display of the instruction of the next trial. Each session consisted of eight blocks of 4:52 minutes of NFB training.

### Real-time EEG signal processing and feature extraction

All signal processing and feature extraction were implemented in MATLAB (version R2022b, MathWorks Inc., USA). The NFB system collected EEG data into 250-sample windows (250 ms), every 100 ms (i.e., with an overlap of 150 samples, 150 ms), hence resulting in feedback updates via the arrowhead every 100 ms. The DC offset of the data for each channel within each window was first removed before applying an adaptive filter (He et al., 2004) to remove the eye-muscle-related activity. The adaptive filter used the vertical (VEOG) and horizontal (HEOG) electrooculogram channels as references. The signals were then filtered with a spatial Laplacian filter (Perrin et al., 1989), followed by the calculation of the Power Spectral Density (PSD) using the Discrete Fourier Transform (fft function) in MATLAB. Specifically, each data window was zero-padded to a length of 1000 samples. Spectral power was averaged across relevant electrodes and frequencies. Table I shows the electrode locations, according to the extended 10-20 electrode placement standard, and frequency range for each feature. The averaged power was then used to calculate the feedback provided to the participants (direction and pointiness of arrowhead; calculated as the ratio between this power measure and the threshold).

### Offline signal processing

The following steps were performed in MATLAB R2022b using custom scripts that incorporated functions from the EEGLAB toolbox (Delorme & Makeig, 2004). The raw EEG data from each participant were imported into EEGLAB. The data was then downsampled from 1000 Hz to 100 Hz to reduce computational load. A bandpass filter between 1 and 40 Hz was applied using pop_eegfiltnew with a Hamming windowed sinc FIR filter to remove slow drifts and high-frequency noise, including line noise. In a first pass of signal processing, we cleaned and removed noisy data for decompositions using Independent Component Analysis (ICA). The clean_rawdata function, an EEGLAB plugin implementing the Artifact Subspace Reconstruction (ASR) method (Mullen et al., 2015), was used for preliminary artifact removal. The following parameters were used: FlatlineCriterion: 40, ChannelCriterion: -1, LineNoiseCriterion: 0.85, BurstCriterion: 4, WindowCriterion: 20, BurstRejection: 0.25 and Distance: ‘Euclidian’ to identify and remove bad channels and high-amplitude artifacts. If ASR removed channels in the previous step, they were interpolated using spherical spline interpolation (pop_interp function) based on the remaining good channels (Perrin et al., 1989). The data was re-referenced to the average of all channels to provide a common reference for subsequent analysis. ICA was performed using the PICARD algorithm (pop_runica function), an optimized Infomax ICA algorithm for detecting source non-Gaussianity. The current study selected PICARD because it is optimized for high-density EEG recordings. The number of components extracted was determined by principal component analysis, retaining only components with eigenvalues larger than 1e-7 (a threshold set by the PICARD algorithm), ensuring no relevant information was discarded (Hyvärinen & Oja, 2000). The resulting independent components (ICs) were automatically classified using the ICLabel algorithm (Pion-Tonachini et al., 2019). ICLabel assigns probabilities to each IC belonging to different categories (e.g., brain, muscle, eye, channel noise). ICs with a probability greater than 0.8 belonging to the “Eye” category were marked for rejection.

In the second pass, in which the resulting data were used for subsequent analyses, we used the raw EEG data at 1000 Hz, which was bandpass filtered using the same settings as in the previous steps. Critically, in the second pass of the clean_rawdata function, we did not remove artifactual data; noisy portions were instead reconstructed. In this step, the following parameters were used to clean significant artifacts, using a BurstCriterion: 20 and not rejecting any channels or time segments based on criteria other than the burst criterion: FlatlineCriterion: ‘off’, ChannelCriterion: ‘off’, LineNoiseCriterion: ‘off’, WindowCriterion: ‘off’, BurstRejection: ‘off’, Distance: ‘Euclidian’. The ICs identified as artifacts (eye activity) from the first pass were subtracted from the data using the pop_subcomp function. This step effectively removed the identified artifacts from the EEG data (Makeig et al., 1996).

A surface Laplacian filter was applied to the cleaned EEG data using the Perrin et al. (1989) algorithm to attenuate the effects of volume conduction. The regularization parameter (lambda) was set to 1e-5, and no smoothing constant was used.

### Offline feature extraction

Time-frequency decomposition was performed using a complex Morlet wavelet transform applied to the recording. The parameters for the wavelet analysis were as follows: number of frequencies: 40, linearly spaced from 1 to 40 Hz; number of cycles: fixed at 7 across all frequencies; and wavelet time: -2 to +2 seconds. Following time-frequency decomposition, the neurofeedback task-related data segments were extracted from each channel and each frequency. The power of the EEG signal at each time-frequency point was calculated by taking the squared magnitude of the complex time-frequency coefficients obtained from the wavelet convolution. The power values were converted to decibels (dB) using the formula 10 * log10(power/mean (power)), where the average power was calculated across the whole recording for each frequency and channel. This dB normalization is commonly used in EEG analysis to account for individual differences in overall power and highlight relative power changes over time. We retained samples only during the NFB training epochs of the task. The power values were extracted at every 10th sample from the time-frequency representation (corresponding to a down-sampling to 100 Hz). Finally, the power values were averaged across the NFB feature-specific channels and frequency band for NFB training for each participant.

### Offline statistical analysis Fitting learning curves

Following offline preprocessing and feature extraction, the EEG data was analyzed using penalized B-splines, implemented in R (version 4.2.1; R Core Team, 2012). This analysis aimed to model the trajectory of EEG power (in dB) over the course of each neurofeedback session for each participant and for each feature separately. To model the non-linear changes in EEG power over time, penalized B-spline basis functions were created using the bbase function from the JOPS package (version 0.1.2; Eilers, 2023). The following parameters were used for constructing the B-spline basis: time vector (in minutes) ranging from the start to the end of the NFB session, minimum value of the time vector, and maximum value of the time vector. 127 equally spaced segments and 3 degrees of the B-splines, cubic B-splines. The number of segments and degrees determines the number of B-spline basis functions, calculated as segments + degrees. We selected 127 basis functions to place the “peak” of the basis functions approximately at the start, middle, and end of each of the 64 trials, which were 30 seconds each. A penalized regression model was fitted to the data for each participant and each feature separately. The model can be represented as:

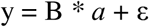

where y is the vector of EEG power values (in dB), B is the B-spline basis matrix, *a* is the vector of unknown coefficients, and ε is the error term. A penalty term was introduced based on the differences between adjacent B-spline coefficients to prevent overfitting and ensure a smooth fit. The penalty term is defined as:

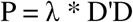

Where λ is the smoothing parameter that controls the trade-off between fidelity to the data and the smoothness of the fit, and D is a difference matrix that computes the third-order differences between adjacent coefficients (a common choice for cubic B-splines to penalize the integrated squared second derivative, which measures the curvature of the function, see e.g. Wood, 2017). The penalized least squares solution for the coefficient vector *a* is obtained by minimizing the following objective function:

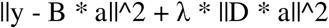

which leads to the following closed-form solution: a = (B’B + λ * D’D)^^-1^ * B’y

The optimal smoothing parameter λ was determined using the V-curve method (Frasso & Eilers, 2014). This method involves fitting the model with a series of λ values (ranging from 10^^2^ to 10^^7^ in this case, with a step size of 0.1 on the log10 scale) and plotting the corresponding values of the residual sum of squares (fit) and the penalty term (roughness) on a log-log scale. The optimal λ corresponds to the V-curve’s corner, representing the best balance between fit and smoothness. The V-curve was calculated by computing the Euclidean distance between adjacent points on the curve and selecting the λ corresponding to the minimum distance. The penalized regression model was fitted to the data for each participant using the optimal λ value determined in the previous step. The estimated coefficient vector and the fitted values of EEG power (in dB) were computed and saved for further analysis.

Bayesian standard errors were calculated for the smooth fit using the method described by (Wand, 2003), providing a measure of uncertainty around the estimated smooth trajectory. This involved computing the variance-covariance matrix of the fitted values and deriving the standard deviation for each time point.

### Identification of Learners

To identify participants who successfully modulated their EEG activity during the NFB training, we calculated the proportion of time points where the upper (for up-regulated features)/lower (for down-regulated features) bound of the Bayesian confidence interval (calculated as the fitted value ± three times the standard error) corresponding to 99.7% confidence of the fitted EEG power was significantly above (for up-regulated features: Theta and Alpha) or below (for down-regulated features: SMR and Beta) the individual’s baseline threshold. Only time points after the initial four minutes (the baseline period) were considered. In simple terms, this approach quantified how often a participant’s EEG power was reliably above or below the target threshold during the NFB session. Participants were classified as *“learners”* if they maintained the target EEG activity for more than 50% of the training time following the baseline period. In addition to the proportion of time, we also extracted two additional metrics: 1) the mean amplitude of the fitted power across all time points after the baseline period, and 2) the length of the longest continuous run of time points where the power was either above or below the threshold (depending on the regulated feature) expressed as a percentage of the total number of time points after the baseline. This “run length” metric measures the consistency of successful modulation.

### Clustering of Learners

To identify subgroups of learners with distinct temporal profiles of EEG power during the NFB sessions, a time-series clustering analysis was performed using the k-shape algorithm (Paparrizos & Gravano, 2015), implemented in the dtwclust package (version 5.5.11; Sardá-Espinosa, 2019) in R. The clustering analysis was based on the coefficients of the penalized B-spline models fitted to each participant’s EEG power trajectory (as described in the previous steps). Inspired by Iorio et al. (2016), this approach leverages the dimensionality reduction achieved by the spline basis expansion, using the coefficients as a parsimonious representation of the time series while retaining the essential information about its shape. The coefficient vectors combined for each feature (Theta, Alpha, SMR, Beta) were standardized using z-score normalization to ensure that all coefficients contributed equally to the cluster distance calculations, regardless of their original scale. The resulting matrix was then converted into a time-series list object (using the tslist function). The k-shape algorithm was applied to the time-series list using the tsclust function. Two clusters were predefined based on prior visual inspection of each participant’s fitted EEG power trajectory. We used a shape-based distance measure specifically designed for time-series data that is invariant to scaling and shifting (Paparrizos & Gravano, 2015). The k-shape algorithm was repeated 1000 times with different random initializations to mitigate the influence of the initial centroid selection on the final clustering solution. To obtain a robust clustering solution, an ensemble clustering approach was employed. The resulting 1000 k-shape solutions were then combined into a consensus clustering using the cl_medoid function (clue package, version 0.3-65; Hornik, 2005). The cl_medoid function identifies the most representative clustering solution from the ensemble (the medoid) that minimizes the average distance to all other solutions in the ensemble. The resulting consensus clustering was used as the final clustering solution. The distance between each learner and the medoid of their assigned cluster was also computed (using the cldist function) as a measure of the quality of the cluster assignment. Given the limited sample size, particularly for non-learner groups, the clustering results should be interpreted as exploratory and primarily descriptive of potential learning dynamics.

### Modeling EEG Power with Mixed-Effects Models

To investigate spatial and temporal dynamics of EEG power during the NFB training, and how these dynamics differed between the identified clusters (from the previous clustering analysis), we employed linear mixed-effects models with tensor product interactions. These models were fitted using the bam function from the mgcv package (version 1.9-1; Wood, 2025) in R. The bam function is specifically designed for large datasets and uses optimized algorithms for efficient model fitting.

To visualize the average spatiotemporal dynamics of EEG power for each identified cluster, while accounting for individual differences, we fitted a generalized additive mixed model (GAMM). The analysis focused on the B-spline coefficients derived previously, representing EEG power over time for each participant and electrode.

### Data Preparation

The B-spline coefficients were combined into a long-format data frame containing the participant ID, electrode label, coefficient number (indexing time within the session, 1-128), and the coefficient value (amplitude). Cluster assignments obtained from the prior analysis were merged into this data frame. The spatial locations (Cartesian coordinates Cartesian_X, Cartesian_Y) for each electrode were added using the electrode_locations function from the eegUtils package (version 0.4.6; Craddock, 2024), based on a standard extended 10-20 system layout. An interaction variable (clusterChannelInteraction) was also created to represent the combination of cluster assignment and whether an electrode belonged to the predefined region of interest (ROI) for the specific EEG feature being analyzed.

### Model Specification

We specified a GAMM using the bam function from the mgcv package (version 1.9-1, Wood, 2017), designed to capture the key sources of variation in the data. The model structure was chosen to allow for distinct spatiotemporal patterns across clusters and to account for participant-level variability:

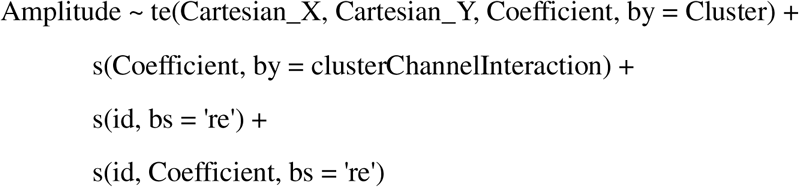

The dependent variable was amplitude (the B-spline coefficient value). The tensor product smooth te(Cartesian_X, Cartesian_Y, coeff, by = Cluster) captures the primary interaction of interest, modeling how the smooth pattern of EEG power across space (Cartesian_X, Cartesian_Y) and time (Coefficient) differs between the cluster groups. The term s(Coefficient, by = clusterChannelInteraction) models the overall time course (Coefficient) separately for each combination of cluster and ROI assignment (clusterChannelInteraction), allowing the average temporal trend within the ROI to differ by cluster. The random intercepts for participant id s(id, bs = ‘re’) account for overall individual baseline differences. The random slope s(id, Coefficient, bs = ‘re’) for coeff varying by id allows the individual time courses to deviate from the group average trend. These random effects are crucial for appropriately modeling the inherent hierarchical structure of the data (participants measured repeatedly over time and space).

### Model Application for Visualization

While alternative model structures could be conceived, this specification was deemed sufficient and theoretically appropriate for the primary goal: generating interpretable visualizations of the central tendencies and variability within each cluster. Formal model comparison using AIC was briefly explored as a sanity check, but the emphasis was not on selecting the single “best” statistical model in an AIC sense. Instead, diagnostic checks (e.g., Q-Q plots, residual distributions) were performed on the chosen model to ensure it reasonably represented the data structure without evident violations of statistical assumptions (Wood, 2017).

### Generating Visualizations

The fitted model was then used to predict each cluster’s expected amplitude across all electrodes and time points. These predictions form the basis for the visualizations. Averaging predictions within specific time blocks allowed for the creation of topographical plots illustrating the estimated spatial distribution of EEG power for each cluster throughout the session. Averaging predictions across electrodes within the ROI (or globally) generated the estimated average time course of EEG power for each cluster. Confidence intervals around these trend lines were computed from the model’s posterior distribution. These intervals primarily illustrate the uncertainty in the estimated average pattern and provide a visual sense of the variability within each cluster rather than making strict statistical inferences at every single time point (Simpson, 2024; Wood, 2017).

### Qualitative Analysis of Self-Reported Strategies

To systematically analyze the qualitative self-report data, a thematic analysis was conducted using the large language model Gemini 2.5 Pro (Google AI) on de-identified data. All personal identifiers were removed prior to analysis, and the dataset contained only non-identifiable textual responses alongside study-specific variables (participant ID codes, target EEG feature, and cluster assignment). The data were compiled into a structured format (CSV) for analysis.

The analysis followed an iterative prompting process. First, the model was provided with the complete dataset and tasked with identifying emergent themes in the strategies used for each EEG feature. Subsequent prompts directed the model to synthesize themes within each performance cluster and to explicitly compare and contrast the strategies reported by successful learners (clusters 1 and 2) with those of non-learners. This AI-assisted approach was used to aid in efficient identification and synthesis of patterns across the large corpus of text data. A manual, deliberate human analysis of the findings followed this.

### AI Use Disclosure

Generative artificial intelligence (AI) tools were used in a limited and supportive role during the preparation of this manuscript. Specifically, AI was used for (i) language editing and grammatical refinement, (ii) drafting and structuring selected text passages, (iii) assisting in the generation and debugging of analysis code, and (iv) supporting the synthesis of qualitative data during the thematic analysis of self-reported strategies.

All AI-assisted outputs were critically reviewed, edited, and validated by the authors. No results, interpretations, or conclusions were generated without human oversight. The authors take full responsibility for the accuracy, integrity, and originality of all content presented in the manuscript.

## Results

### Overall Performance of Neurofeedback Training

We first summarize the participant’s ability to modulate the four target EEG features across all sessions. As shown in Figure 2, the participants exhibited considerable variability in their ability to control the different EEG features. Notably, none of the participants could successfully up-regulate frontal Theta activity, as indicated by the uniformly red cells in the Theta row (0% success rate across all participants). In contrast, Alpha up-regulation was generally more successful, with most of the participants (60%) being able to increase Alpha power over the threshold during at least 50% of the session. Down-regulation of SMR and Beta activity showed even more learners. Only a few participants were unable to control the SMR or Beta successfully. Overall, 95% of the participants could down-regulate SMR, and 90% down-regulated Beta.

**Figure 2.**
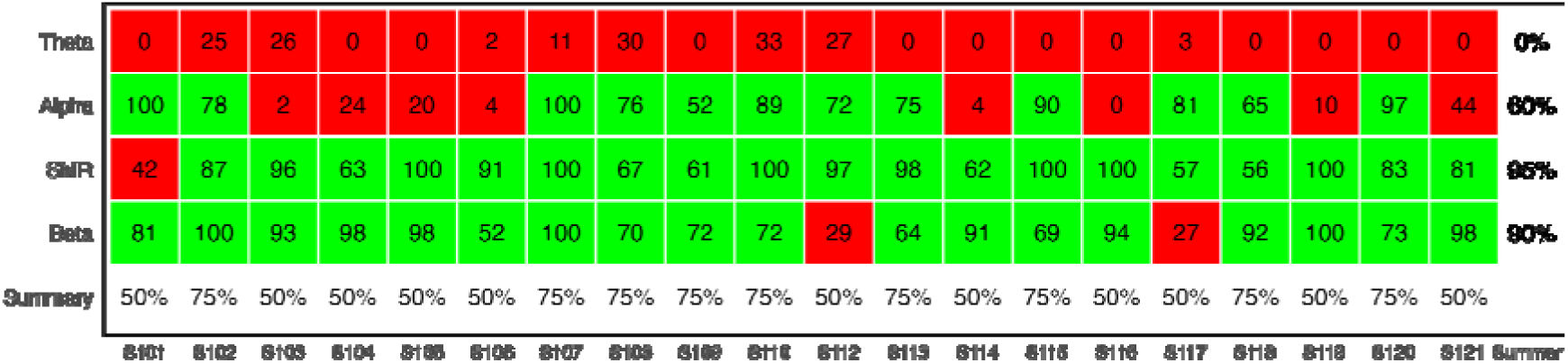
Binary heatmap summarizing the percentage of time each participant successfully modulated each EEG feature. Green cells indicate successful modulation for at least 50% of the session; red cells indicate unsuccessful modulation (<50% of the session). The bottom row shows the percentage of features each participant successfully controlled. The rightmost column shows the percentage of participants who successfully controlled each feature.

Notably, all participants (100%) could control at least two out of the four features in one single NFB session, while 45% (9 out of 20) could control three features. These results suggest that while there is substantial inter-individual variability in NFB performance, all participants could achieve control over part of their EEG activity.

### The dynamics of Neurofeedback Training

Next, we characterize the temporal and spatial EEG activity dynamics for each feature with topographic plots and trendlines across time for non-learners and the two clusters of learners. Notably, given the absence of successful theta regulation, clustering in this feature was used to characterize inter-individual differences in EEG dynamics among non-learners.

### Performance in up-regulating frontal midline Theta

As evident from Figure 2, none of the participants could learn to self-regulate frontal midline Theta within a single session. However, we performed a cluster analysis to explore whether we could see differences on the group level in non-learners. In the small subgroup Cluster 1 (n= 3 participants), we observed a relative parietal Theta positivity that decreased during the session, extending forward in midline regions. The predicted activity in frontal midline channels varied throughout the session, with a sharp decrease during block 8 at the end of the session. For Cluster 2 (n=17 participants), we observed a high global relative positivity at the start of the session. Similarly, compared with Cluster 1, this positivity decreased throughout the session, with the highest power reduction in midline frontal regions.

Although the evolution of Theta power activity was opposite to what was expected, it was spatially localized to the expected locations. Next, we investigate the frequency specificity of Theta evolution across the session. By plotting the difference between the 8^th^ block and the baseline for each frequency band across frontal midline electrodes, as Figure 3 suggests, Theta decreased relative to the threshold in the last block for both clusters (Figure 4; red and blue violin plots for Theta; 95% bootstrapped CI<0, p<0.05). Figure 4 shows that this decrease was specific to the Theta band in Cluster 1. However, the reduction in power for Cluster 2 was accompanied by a decrease in all frequency bands (Alpha, SMR, and Beta; 95% bootstrapped CI<0, p<0.05).

**Figure 3.**
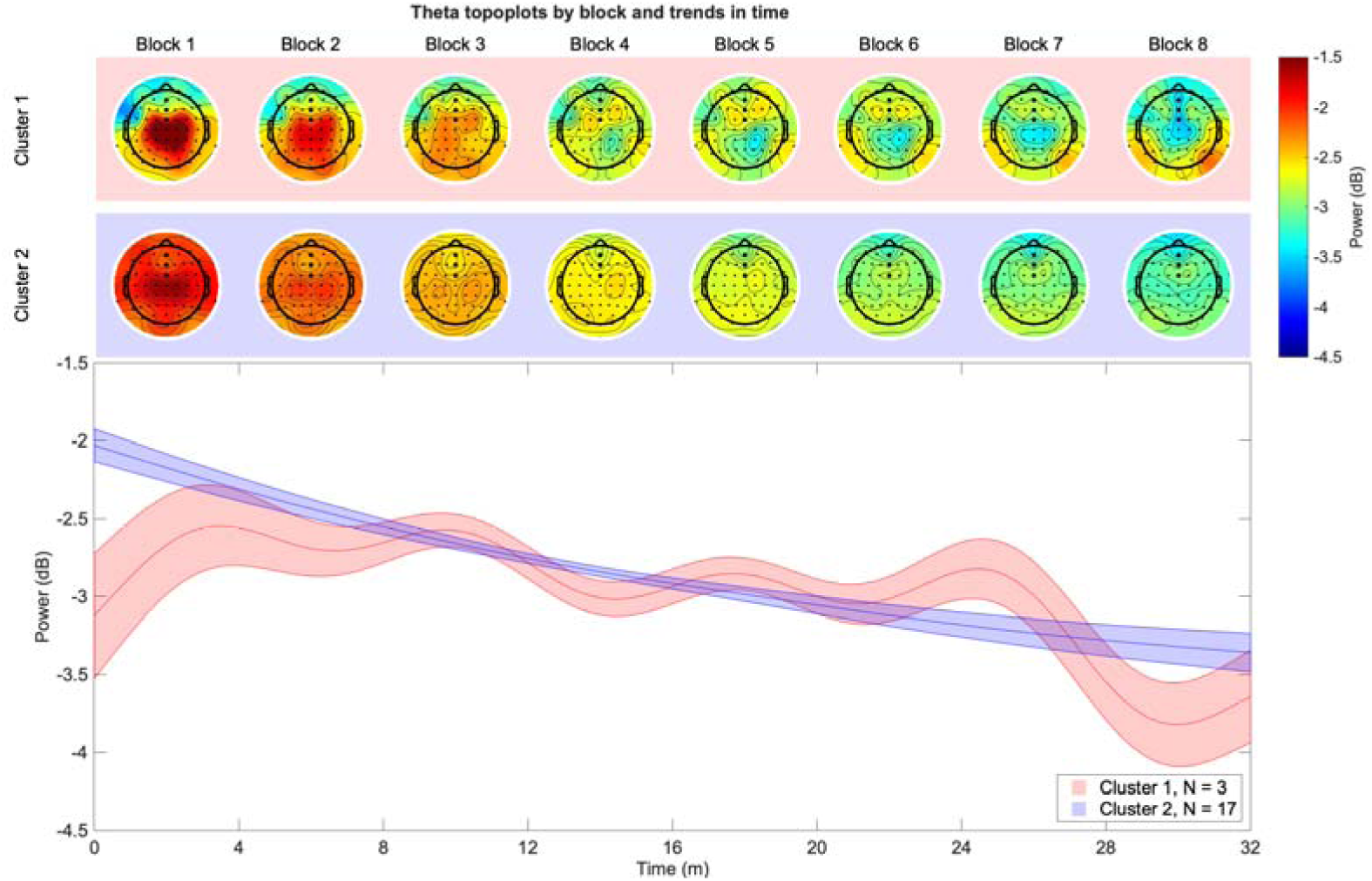
Top two rows: Topographic plots with average activity in eight blocks of frontal midline Theta neurofeedback by cluster. Bottom panel: Trendlines (with 95% CIs) show the predicted activity in each cluster across the recording. Note that both clusters in this plot were non-learners.

**Figure 4.**
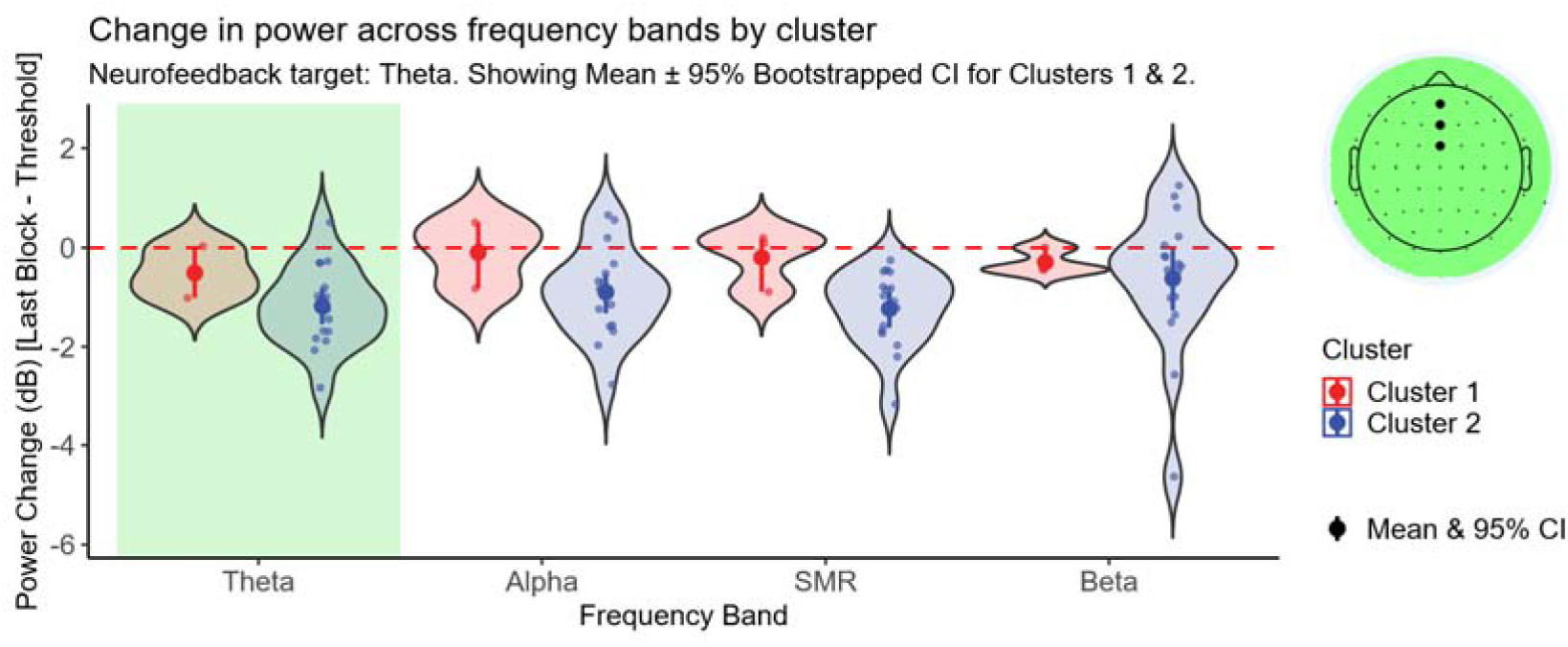
Frontal midline Theta change (last block - threshold) for all clusters in each frequency band. Spread and central tendency of the change in EEG power (dB), calculated as the average power in the last block minus the baseline threshold power, across different frequency bands. Data are shown separately for participants assigned to Cluster 1 (red) and Cluster 2 (blue). The visualization combines violin plots (showing the probability density of the data), jittered individual data points, and point-range plots indicating the group mean and its 95% bootstrapped confidence interval. A dashed red line at 0 dB indicates the point of no change relative to the threshold. The target frequency band for the neurofeedback training analyzed here (Theta) is highlighted with a green background rectangle. Confidence intervals not containing zero suggest a statistically significant change from baseline at the alpha = 0.05 level.

### Performance in up-regulating occipital Alpha

Compared to frontal midline Theta, the participants were more successful in learning to self-regulate occipital Alpha. For Cluster 1 (n=7 participants), we observed an increase bilaterally across occipital channels. The increase continued throughout the session (cf. Figure 5). For Cluster 2 (n=5 participants), we observed a tendency for a more right-lateralized occipital increase that plateaued following block 3. A reduction in Alpha over bilateral centro-parietal locations (motor areas) could be observed during the last blocks of the session (from session 4 to 5). For non-learners (n=8 participants), we observed a continuous decrease in Alpha.

**Figure 5.**
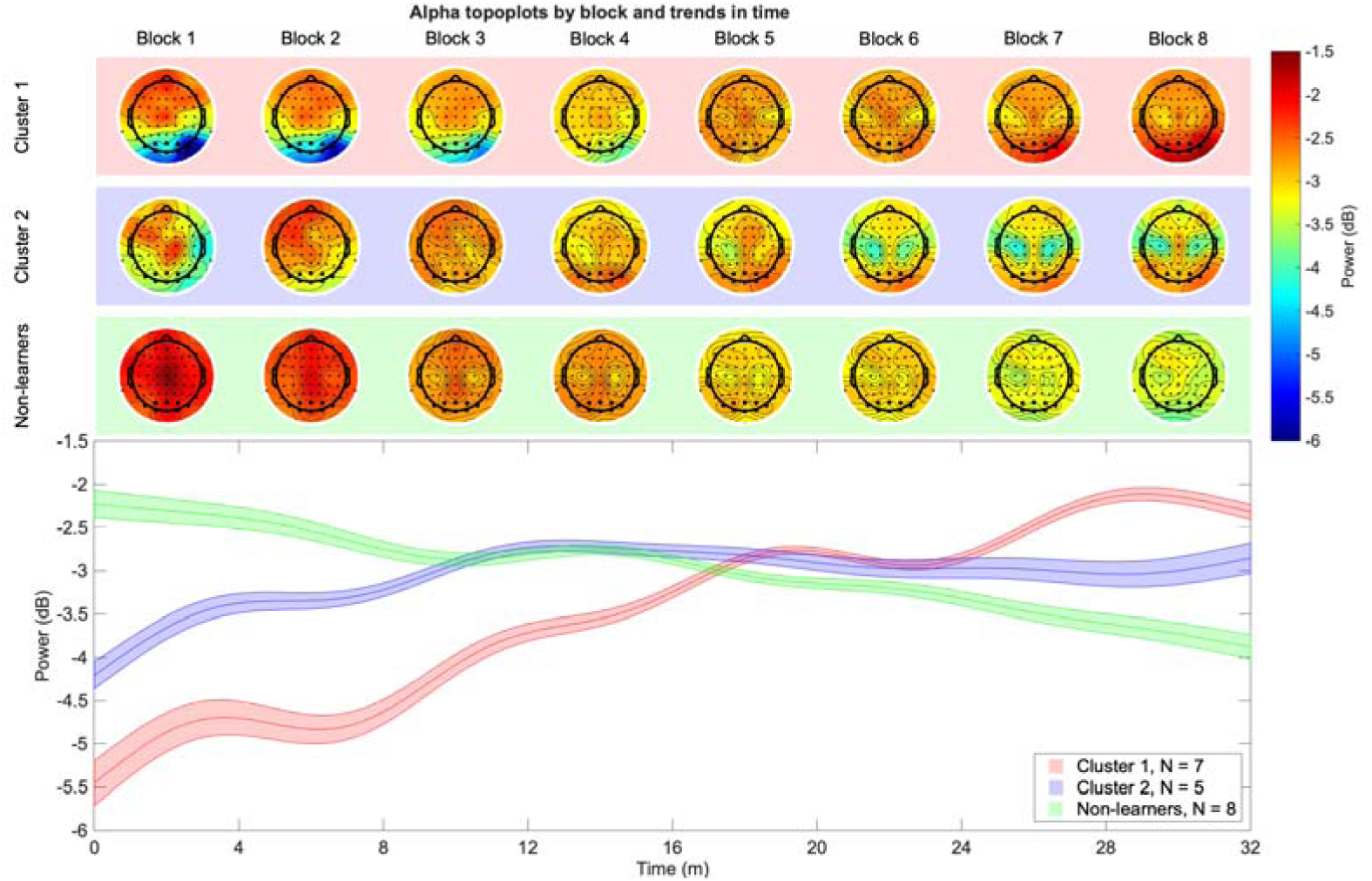
Top three rows: Topographic plots with average activity in eight blocks of occipital Alpha neurofeedback by cluster, with the last row showing non-learners. Bottom panel: Trendlines (with 95% CIs) show the predicted activity in each cluster across the recording. Note that the cluster colored in green was non-learners.

In the frequency domain, we observed the largest positive change (last block - baseline) for Cluster 1 in the Alpha band with slight increases in the SMR and Beta bands, but not the Theta band (Figure 6; red violin plots; 95% bootstrapped CI>0, p<0.05). For Cluster 2, the increased power in Alpha was not associated with increases in other frequency bands, however a significant decrease in the Theta band was observed (Figure 6; blue violin plot for Theta; 95% bootstrapped CI<0, p<0.05). For the non-learners, we observed a decrease in power across all frequency bands (Figure 6; green violin plots; 95% bootstrapped CI<0, p<0.05).

**Figure 6.**
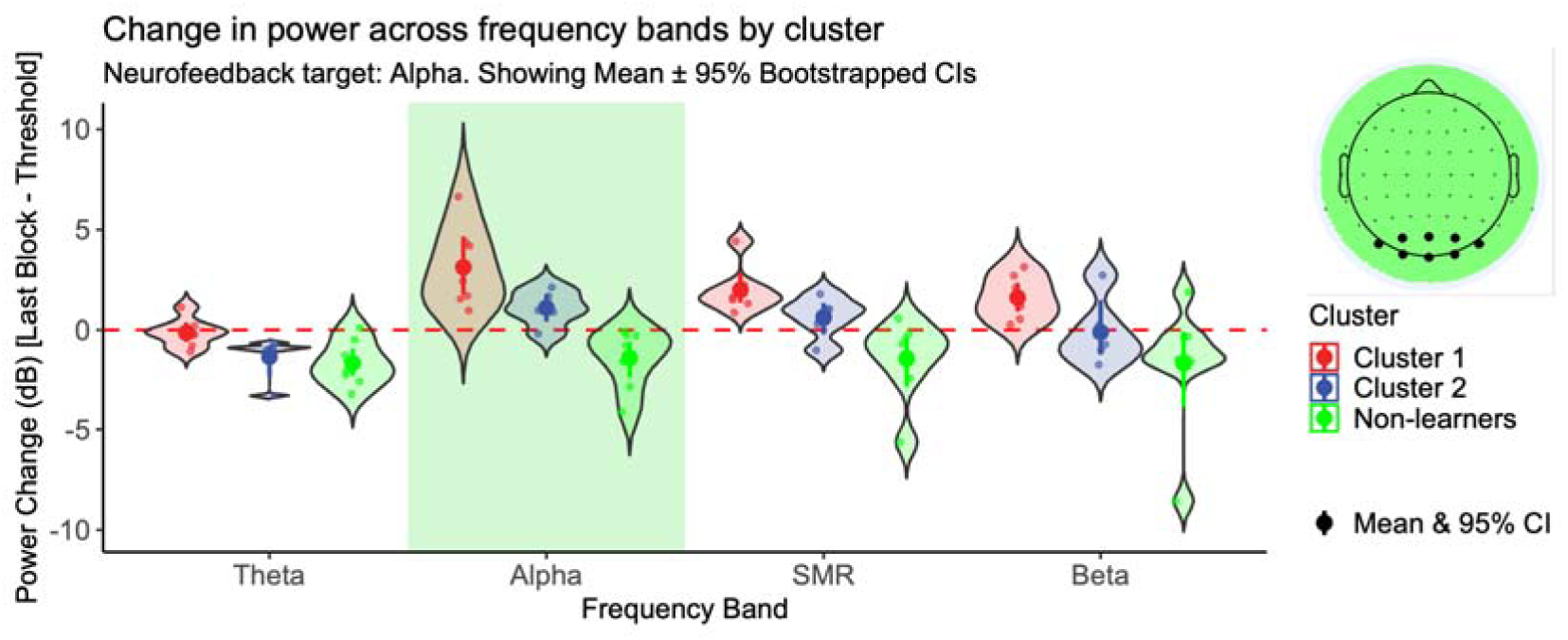
Occipital Alpha change (last block - threshold) for all clusters in each frequency band. Spread and central tendency of the change in EEG power (dB), calculated as the average power in the last block minus the baseline threshold power, across different frequency bands. Data are shown separately for participants assigned to Cluster 1 (red), Cluster 2 (blue), and non-learners (green). The visualization combines violin plots (showing the probability density of the data), jittered individual data points, and point-range plots indicating the group mean and its 95% bootstrapped confidence interval. A dashed red line at 0 dB indicates the point of no change relative to the threshold. The target frequency band for the neurofeedback training analyzed here (Alpha) is highlighted with a green background rectangle. Confidence intervals not containing zero suggest a statistically significant change from baseline at the alpha = 0.05 level.

### Performance in down-regulating unilateral centrotemporal SMR

Figure 7 illustrates the spatiotemporal dynamics of SMR power for the identified learners’ clusters and the non-learners. For Cluster 1 (n=7 participants), we observed an initial bilateral decrease in SMR power spanning widespread centro-parietal and occipital regions. This decrease evolved into a more focused, left-lateralized reduction primarily in left centro-parietal electrodes. This pattern persisted until approximately block 4, after which the activity plateaued. Cluster 2 (n=12 participants) exhibited a consistent left-lateralized decrease in SMR power, concentrated over left centro-parietal areas. The SMR activity across the feature channels decreased steadily throughout the session, slightly plateauing during the last block. The non-learner, consisting of a single participant, presented a different pattern: SMR power in the self-regulated channels remained without noticeable change until Block 6, and from that point, there was a decrease in power until the end of the recording. As shown in the lower panel of Figure 7, the overall trend throughout the session shows a decreasing amplitude for Cluster 1 and Cluster 2 and a stable amplitude for the non-learner until the later part of the recording. Cluster 1 showed a steeper decrease than Cluster 2, and the change in Cluster 2 plateaued around minute 16.

**Figure 7.**
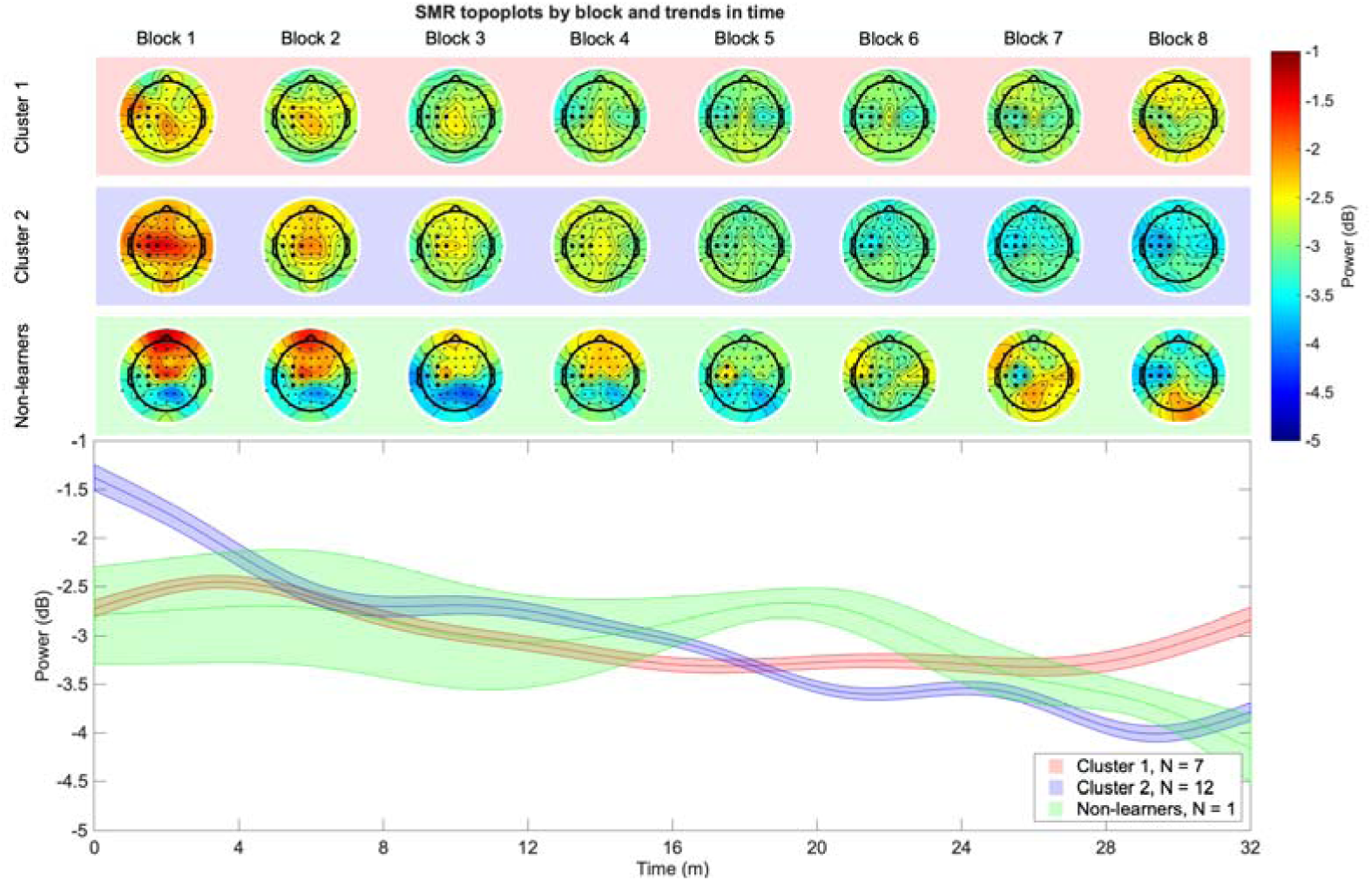
Top three rows: Topographic plots with average activity in eight blocks of centrotemporal SMR neurofeedback by cluster, with the last row showing non-learners. Bottom panel: Trendlines (with 95% CIs) show the predicted activity in each cluster across the recording. Note that the cluster colored in green was non-learners. Note: The electrode positions for two left-handed participants were flipped across the midline in the topographic plots only for visualization purposes. All quantitative analyses, including modelling and clustering, were performed on the original (non-flipped) data, to preserve lateralized information.

In the frequency domain, SMR down-regulation did not show frequency specificity. Figure 8 displays the change in power (in dB) between the last block of the NFB session and the individual’s baseline threshold, broken down by frequency band (Theta, Alpha, SMR, Beta) and cluster (Cluster 1, Cluster 2, Non-learners). The boxplots illustrate the distribution of these power changes across participants within each group.

**Figure 8.**
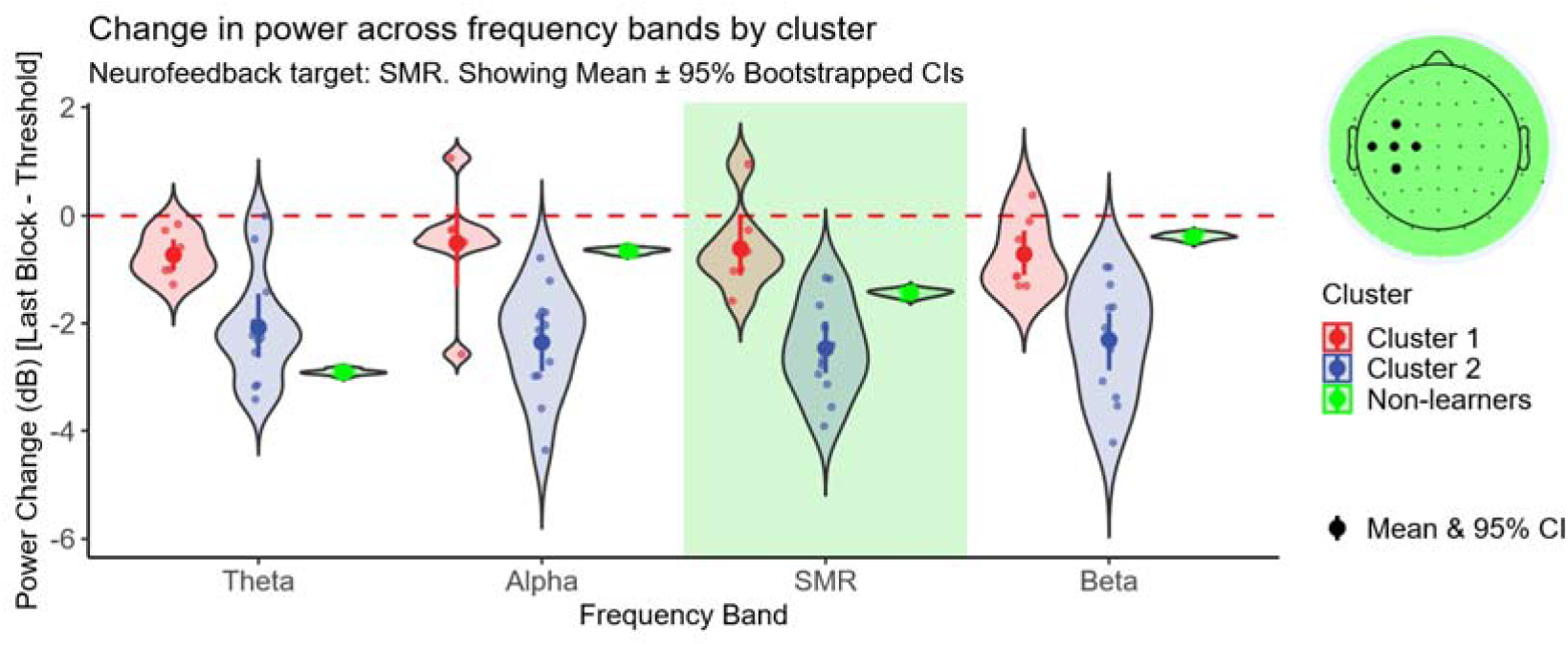
Centrotemporal SMR change (last block - threshold) for all clusters in each frequency band. Spread and central tendency of the change in EEG power (dB), calculated as the average power in the last block minus the baseline threshold power, across different frequency bands. Data are shown separately for participants assigned to Cluster 1 (red), Cluster 2 (blue), and non-learners (green). The visualization combines violin plots (showing the probability density of the data), jittered individual data points, and point-range plots indicating the group mean and its 95% bootstrapped confidence interval. Due to the small sample size in the non-learner group, data points were oversampled with negligible jitter to generate the violin density shape. A dashed red line at 0 dB indicates the point of no change relative to the threshold. The target frequency band for the neurofeedback training analyzed here (SMR) is highlighted with a green background rectangle. Confidence intervals that do not contain zero suggest a statistically significant change from baseline at the alpha = 0.05 level.

Both clusters 1 and 2 learners show a decrease in SMR power (Figure 8; red and blue violin plots for SMR; 95% bootstrapped, CI<0, p<0.05), but the magnitude of this decrease differs substantially. Cluster 2 exhibits a notably larger decrease in SMR power compared to Cluster 1. A similar decrease in power was evident in all the other frequency bands (Figure 8; red and blue violin plots; 95% bootstrapped, CI<0, p<0.05). Although we see a reduction in SMR power for the non-learner, this activity was not sustained long enough to reach the defined threshold of 50% of time above or below the threshold for successful neural self-regulation.

### Performance in down-regulating central Beta

Figure 9 illustrates the spatiotemporal dynamics of Beta power for the identified learners’ clusters and the non-learners. In the very small subgroup, Cluster 1 (n=3 participants), the initial blocks show a widespread decrease, most prominent over central electrodes; this pattern appears to be largely driven by an emerging decrease in Beta in frontal and more peripheral and posterior areas of the scalp. This central decrease intensified between blocks 3 and 5, followed by a stabilization or plateau from block 6 onwards. Cluster 2 (n=15 participants) exhibited a more localized decrease from the start of the NFB session. Initially, the topographic maps show a circumscribed reduction in Beta power centered over central electrodes. From block five and throughout the remainder of the session, the decrease seems to be driven by a left-lateralized centro-parietal decrease in Beta. The time course representation, depicted in the lower panel, shows a steep decrease in Beta power for Cluster 2, and a more gradual decrease in amplitude for Cluster 1. Cluster 1 seemed to plateau around 16 minutes, whereas the decrease for Cluster 2 remained until the last block. The non-learners (n=2 participants) showed a similar spatial pattern to Cluster 1 but started from a lower amplitude and maintained a near-constant level of Beta amplitude throughout the session. However, the results should be treated cautiously due to small sample sizes in the Cluster 1 and non-learners.

**Figure 9.**
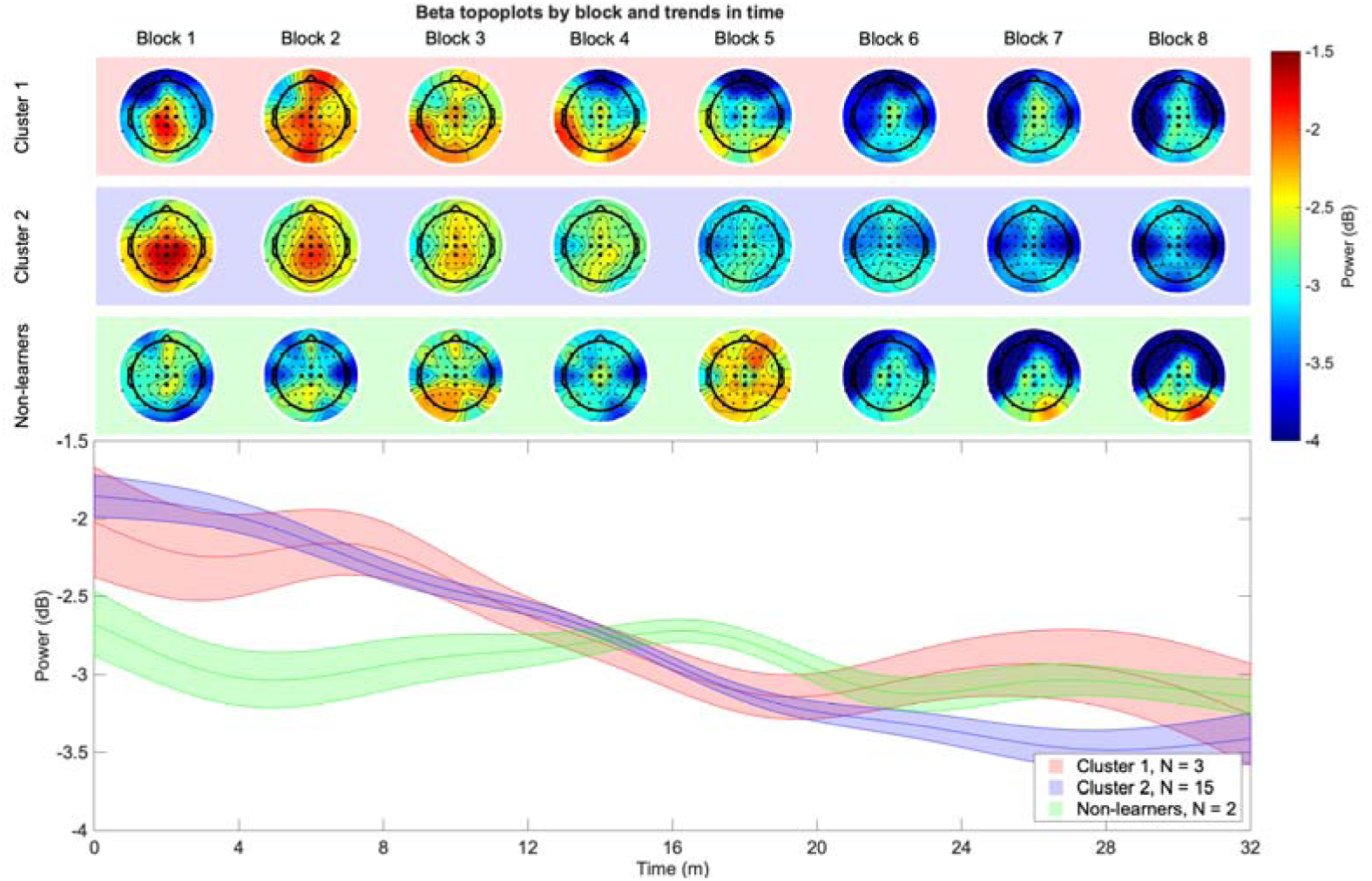
Top three rows: Topographic plots with average activity in eight blocks of central Beta neurofeedback by cluster, with the last row showing non-learners. Bottom panel: Trendlines (with 95% CIs) show the predicted activity in each cluster across the recording. Note that the cluster colored in green was non-learners.

Similar to SMR, we see a decrease in Beta power in both learner clusters, consistent with successful down-regulation, between the last block and the threshold (Figure 10; red and blue violin plots for Beta; 95% bootstrapped, CI<0, p<0.05), but the magnitude of this decrease differs substantially between clusters. Cluster 2 exhibits a notably larger decrease in SMR power compared to Cluster 1. The Beta reductions in Clusters 1 and 2 were accompanied by a global reduction in power in all other frequency bands (Theta, Alpha, and SMR, Figure 10; 95% bootstrapped, CI<0, p<0.05). The two non-learners reduced power in all frequency bands but the Beta band (Figure 10; green violin plots; 95% bootstrapped, CI<0, p<0.05).

**Figure 10.**
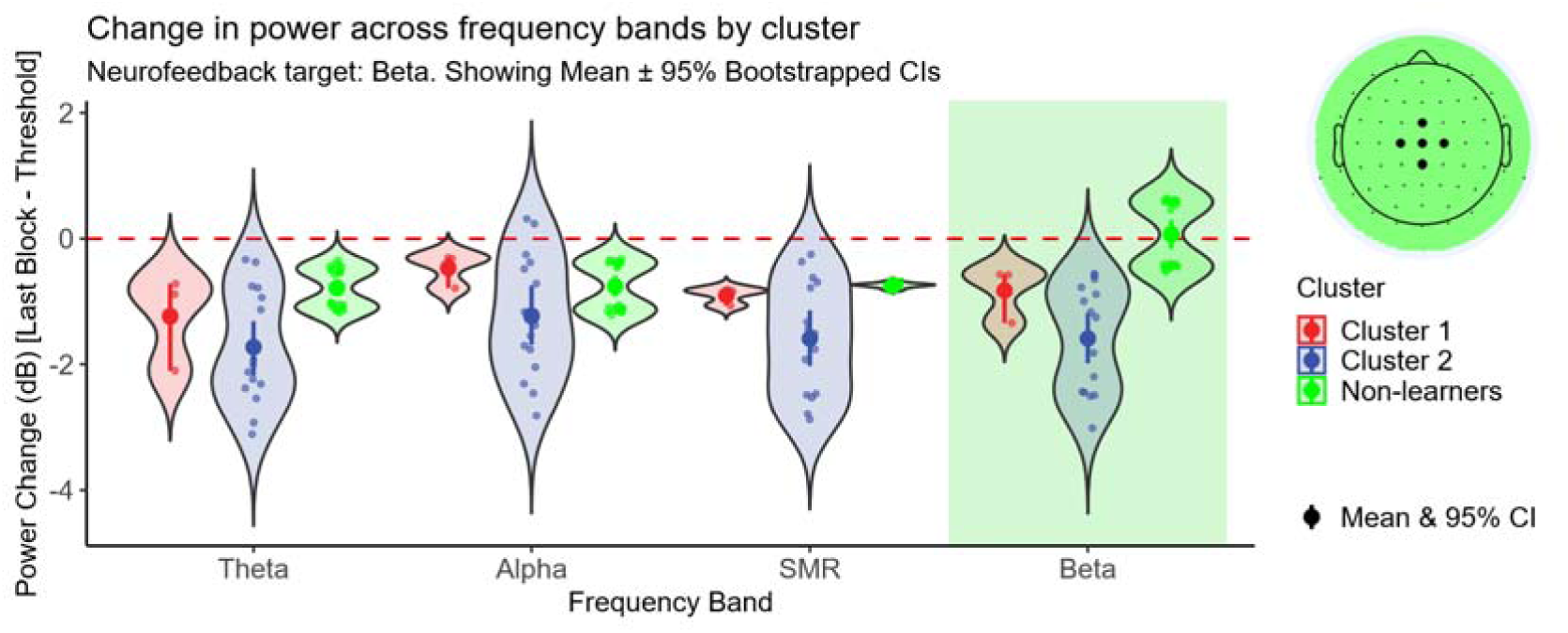
Central Beta change (last block - threshold) for all clusters in each frequency band. Spread and central tendency of the change in EEG power (dB), calculated as the average power in the last block minus the baseline threshold power, across different frequency bands. Data are shown separately for participants assigned to Cluster 1 (red), Cluster 2 (blue), and non-learners (green). The visualization combines violin plots (showing the probability density of the data), jittered individual data points, and point-range plots indicating the group mean and its 95% bootstrapped confidence interval. Due to the small sample size in the non-learner group, data points were oversampled with negligible jitter to generate the violin density shape. A dashed red line at 0 dB indicates the point of no change relative to the threshold. The target frequency band for the neurofeedback training analyzed here (Beta) is highlighted with a green background rectangle. Confidence intervals that do not contain zero suggest a statistically significant change from baseline at the alpha = 0.05 level.

### Qualitative Analysis of Self-Reported Strategies

The thematic analysis of participants’ self-reported strategies revealed distinct patterns that corresponded with both the target EEG feature and NFB performance.

### General Differences Between Learners and Non-Learners

A clear distinction emerged between successful learners (clusters 1 and 2) and non-learners. Notably, for Theta, the clusters solely distinguished different EEG dynamics within non-learners.

- **Learners** typically described a process of **strategy discovery and stabilization**. They reported actively testing different mental states before identifying one that was effective, which they then attempted to maintain. For example, one successful learner for Alpha reported, “Tried many strategies until s/he found what worked. Visualized a world of gravel, picked up individual grains…”.
- **Non-learners**, in contrast, reported experiences of inefficacy and strategy churn. Their reports were characterized by themes of feeling a lack of control, trying “everything” without success, and being unable to find or sustain a working strategy. A representative report from an Alpha non-learner was, “Tried Everything, didn’t feel s/he could control the arrow.”

*Feature-Specific Strategy Themes (identified by large* language model Gemini 2.5 Pro)

- **Theta Up-regulation:** The strategies reported for Theta were highly varied and reflected a universal struggle. Common themes included vague, effortful commands (”thinking go up”), attempts at relaxation that often led to drowsiness (e.g., “Trying to

not to fall asleep”), and complex cognitive tasks such as counting prime numbers or switching languages. No single effective strategy emerged.

- **Alpha Up-regulation:** Two primary successful pathways were identified:

1. **Cluster 1:** This involves general imagination or visualization or performing a non-visual cognitive task, such as “sorting an array of numbers” or mental arithmetic. Descriptions in this cluster were more elaborate and lengthier compared to Cluster 2.
2. **Cluster 2:** This involved generating physical active scenarios such as bike riding or jumping on a trampoline; one participant reported better performance as getting more tired.
- **SMR and Beta Down-regulation:** Due to the inverted feedback design, strategies reported to point the arrow upwards were those that successfully *down-regulated* SMR and Beta power. The analysis revealed a remarkably consistent theme across almost all successful learners: the use of **active, structured cognitive engagement**. No common themes were found within clusters. Several participants reported being in control; however, they could not explain how. Common strategies included:

- **Sequential Cognitive Tasks:** Counting numbers sequentially (two participants), reciting ordered sequences, or performing mental calculations.
- **Sustained Attentional Focus:** Focusing on a single point on the screen, repeating words or songs internally (two participants), or focusing on curious stories.

For SMR and Beta, only two participants reported using strategies involving motor imagery. First, “Ice skating very smoothly, right foot sliding, and then left, and so on.” And second, relaxing and focusing on their breathing “…exhaling very slowly and inhaling very slowly…”.

## Discussion

This study investigated inter-individual variability in EEG NFB performance across four distinct frequency bands, aiming to shed light on the “non-learner” problem. To our knowledge, no other study has employed an intra-subject multi-session experimental design to investigate the ability of learning to self-regulate different cortical rhythms through NFB. In addition, the selectivity of the regulation in both frequency and spatial domains is studied and compared between the different NFB features, increasing our understanding of possible cortical co-dependences. Our central finding is that the ability to self-regulate neural oscillations is not a subjective trait but is dependent on the specific feature being targeted. We demonstrate that down-regulation of sensorimotor rhythms (SMR and Beta) is achievable for a vast majority of individuals, while up-regulation of occipital Alpha is moderately feasible, and up-regulation of frontal midline Theta was not successfully attained within a single session under the present experimental conditions. Crucially, every participant succeeded in regulating at least two features, challenging the notion of a fixed, universal “non-learner” phenotype. Furthermore, our clustering analysis revealed distinct temporal and topographical learning profiles even among strong learners, suggesting diverse underlying neurophysiological strategies. Specifically, we analyzed two subgroups of temporal learning dynamics and identified one linear-like and one non-linear learning dynamic. An incrementally increasing or decreasing regulation of power has been previously shown across subjects (Belinskaya et al., 2020), however there are no previous reports of differential learning dynamics between subjects. We speculate that these different dynamics are associated with differences in learning rate and effectiveness of mental strategy. Assessing learning during NFB training is crucial to enable adaptation of the feedback to better guide and tailor it to the user.

### Frontal midline Theta

The most striking result was the complete failure of participants to up-regulate fm Theta. Instead of the intended increase, all participants exhibited a decrease in Theta power over the session, which was, interestingly, topographically localized to the targeted frontal midline region. Situating the results in the overall literature is challenging, as frontal midline Theta is typically studied in multi-session designs. However, in the many studies reporting successful fm Theta enhancement across a multi-session intervention (Enriquez-Geppert, Huster, Figge, et al., 2014; Eschmann et al., 2020; Eschmann & Mecklinger, 2022), enhancement rarely occurs during the first sessions. Additionally, although largely unreported, fm Theta has not been observed to increase within sessions, but enhancement rather seems to occur between sessions (Enriquez-Geppert, Huster, Scharfenort, et al., 2014). This may partly explain the absence of up-regulation observed in the present study. Apart from a single session, a specific difference between the present study and many previous studies with successful Theta regulation (Brandmeyer & Delorme, 2020; Enriquez-Geppert, Huster, Scharfenort, et al., 2014; Eschmann & Mecklinger, 2022) is that these studies provided participants with strategies. Although there is currently no consensus within the NFB field on whether to give the participants strategies (Enriquez-Geppert et al., 2017), we speculate that offering instructions to participants might be beneficial for learning outcomes in midline Theta NFB. Further, the results might also be explained by negative feedback loops. As the participants struggle to find an effective strategy, they receive negative feedback. Thus, the observed decrease of Theta power may reflect processes such as error-related negativity; however, this interpretation remains speculative, as the present analyses are based on spectral power measures rather than event-related potentials (ERP), and would require dedicated ERP-based protocols to confirm. The error-related negativity has been reported as a phase resetting of Theta (Trujillo & Allen, 2007). Our clustering results further nuance this finding: one group showed a Theta-specific power decrease, while the other showed a broadband power reduction, perhaps reflecting different strategies, one targeting a specific cognitive state and the other lapsing into general fatigue or disengagement.

The strategy reports of the participants strongly illuminate our finding that none were able to up-regulate Theta. Participants consistently described employing effortful, high-load cognitive strategies (e.g., complex calculations, language switching) or battling drowsiness. Successful strategies for up-regulating frontal midline theta are seldom reported in the literature. However, one report suggests that effective strategies often involve sustained, low-effort attention, mindfulness, or an open-monitoring state of meditation (Brandmeyer & Delorme, 2020). The active, goal-directed striving reported by our participants might have induced a state of high executive load, which is neurophysiologically antithetical to the state of relaxed, focused attention that fosters Theta enhancement. Our results, therefore, highlight a critical pitfall for uninstructed Theta neurofeedback: without guidance towards a state of low-effort focus, participants’ intuitive, high-effort attempts might paradoxically suppress the very rhythm they intend to increase.

A key limitation of the present design is the restriction to a single-session training paradigm. Previous studies indicate that frontal midline Theta enhancement typically emerges across sessions rather than within-session. Therefore, the present findings should not be interpreted as evidence of an inherent inability to regulate fm-Theta, but rather as an indication that such regulation may require extended training, strategy guidance, or consolidation processes. This dissociation suggests that different EEG features may vary not only in overall learnability but also in the timescale over which learning occurs, highlighting the importance of matching neurofeedback protocols to feature-specific learning dynamics.

### Occipital Alpha

Up-regulation of occipital Alpha activity was moderately successful in the present study, with 60% of participants defined as learners. This proportion falls within the range reported in prior Alpha NFB research (Alkoby et al., 2018; Hanslmayr et al., 2005), and our results are supported by findings that alpha up-regulation within single sessions is feasible (Escolano et al., 2014). However, direct comparisons of proportions between studies remain limited by inconsistent definitions of “learner” across studies (Rogala et al., 2016). The link between increased Alpha power and cortical inhibition in the visual system, particularly during relaxed wakefulness, is well established (Klimesch, 1999; Klimesch et al., 2007). Our findings extend this work by revealing distinct spatiotemporal and spectral learning patterns across participants.

A striking observation was that participants, primarily those in Cluster 1, showed the largest sustained training-related gains, with widespread bilateral increases in the occipital region sustained throughout the session (Figure 5, top row). By contrast, Cluster 2 participants exhibited a more right-lateralized Alpha enhancement that plateaued after the third block (Figure 5, middle row). The topographies suggest that learners who engage larger occipital regions have the largest change in occipital alpha, while those with more spatially restricted modulation reached a plateau. Non-learners consistently showed a decrease in Alpha power across the scalp (Figure 5, bottom row); this effect was also visible as a broad band decrease in the occipital channels (Figure 6). Across the targeted electrode locations, alpha showed two different learning dynamics: a linear increase and an initial increase followed by a plateau. Prior research has typically focused on average learning trends across multiple sessions, often reporting a linear increase in alpha power over days (Zoefel et al., 2011). However, characterizing learners by their within-session temporal dynamics has not been previously reported. By utilizing a clustering approach, we revealed distinct learning trajectories that are lost in group-averaged data. In contrast to our results showing distinct linear or plateau-like learning trajectories, a recent study reported that single-session alpha up-regulation followed a U-shaped pattern, suggesting no effective learning occurred within individual sessions (Grosselin et al., 2021). Methodological differences in EEG equipment, target electrode location, or feedback modality might be possible reasons for this inconsistency and highlight the delicate nature of NFB learning within vs. between sessions.

The differences in the magnitude of regulation may reflect the combined influence of trait-like physiological constraints and homeostatic regulation mechanisms. Some reports indicate that an increase in alpha power is not correctly represented as an increase in the magnitude or duration but rather as an enhanced incident rate of alpha spindles (Belinskaya et al., 2020; Ossadtchi et al., 2017). We can speculate that the amplitude of alpha spindles are subject-specific and therefore limits the maximum increase that is feasible for each subject. Alpha power spectra exhibit high test–retest reliability (Napflin et al., 2007) and can be considered as stable individual characteristics. Although the present research investigated the learning trajectories, previous research has shown greater NFB gains in high-baseline Alpha individuals (Naas et al., 2019). Because of our pre-processing procedures, specifically decibel conversion that discards the baseline, we cannot confirm if this is the case in the present study.

Considering non-controlled frequencies (Figure 6), we see that Cluster 1’s Alpha gains were accompanied by modest increases in SMR and Beta, suggesting broader frequency co-modulation. Cluster 2’s Alpha changes were more frequency-specific, in line with previous work (Zoefel et al., 2011), whereas non-learners showed decreases across all bands. Such patterns could reflect differences in neural recruitment strategies, where some learners enhance Alpha through global relaxation states that also affect adjacent bands, while others engage more targeted regulatory mechanisms.

In summary, larger gains in occipital Alpha were associated with a topographically wide occipital Alpha activity (Cluster 1). However, this increase came with a cost when we consider that this increase in occipital Alpha was less specific in the frequency range compared to the more right-lateralized group (Cluster 2), which had a lesser increase in occipital Alpha.

The qualitative analysis suggests distinct cognitive mechanisms driving the performance differences between Alpha clusters. Cluster 1 strategies were characterized by high-load internal attention, involving complex mental arithmetic (e.g., ‘sorting numbers’) or detailed visualization. This aligns with the ‘inhibition-timing hypothesis’ (Klimesch et al., 2007) and findings by Cooper et al. (2003), which suggest that high internal cognitive load generates increased Alpha power to inhibit external sensory intake. In contrast, Cluster 2 participants relied on gross motor imagery (e.g., ‘bike rides’, ‘jumping’). While motor imagery is a powerful modulator of brain activity, it typically induces event-related desynchronization (ERD) in Alpha/Mu bands (Pfurtscheller & Neuper, 2001). This reliance on motor-active strategies may explain why Cluster 2 exhibited an initial increase followed by a plateau, as the desynchronizing nature of motor imagery likely counteracted the goal of Alpha synchronization.

Together, these findings indicate that Alpha up-regulation can be achieved through both suppression of external input and engagement of internal representational systems. Rather than pointing to a single, unitary neural mechanism, the effectiveness of both passive and active strategies underscores the flexibility of Alpha-generating networks and highlights the role of individual differences in selecting and deploying successful regulatory tactics. This contributes to a growing body of evidence suggesting that Alpha-based NFB may not be a monolithic process but rather one that supports multiple, individualized cognitive routes to modulation.

### Centrotemporal SMR and central Beta

The highest success rates were observed for the down-regulation of unilateral centro-parietal SMR (95%) and central Beta (90%). The relative ease of this task likely relates to its more direct link to somatic states. Both SMR and Beta rhythms are suppressed during motor planning, execution, and imagery (McFarland et al., 2000). Participants can leverage relatively concrete mental strategies, such as imagining the sensation of movement (Pfurtscheller et al., 2006). Based on the large body of literature related to the use of NFB to enhance motor-related cortical activity after stroke (Foong et al., 2020; Pichiorri et al., 2011; Prasad et al., 2010), the established “gold standard” strategy for modulating sensorimotor rhythms is motor imagery, the mental rehearsal of movement (Neuper et al., 2005). Remarkably, very few of our successful participants reported using motor imagery. Instead, they consistently converged on active, non-motoric cognitive engagement, such as sequential counting, internal monologue, or complex calculations. These results suggest that control over sensorimotor rhythms can also be achieved through non-motoric mental strategies, albeit perhaps not as frequency-specific as when performing, for example, motor imagery.

On this note, a critical finding in our study was the lack of frequency specificity; successful down-regulation in the SMR and Beta bands was accompanied by a decrease in broadband power across all measured frequencies. This suggests that participants did not achieve fine-grained control over a single rhythm but rather induced a more global state of cortical idling or reduced arousal over the sensorimotor cortex, a mechanism that is effective but not spectrally precise. One possible reason for our non-frequency-specific effects was that our NFB protocol did not penalize activity in surrounding electrodes, nor did it employ relative power of SMR and Beta, but the feedback merely reflected the power amplitude of each frequency band in relation to a baseline (i.e., the first NFB block). There are a few studies, particularly related to attention research, that have employed frequency-selective protocols of up-regulation of central SMR or Beta with simultaneous down-regulation of other frequency bands (Dessy et al., 2020; Liu et al., 2022; Vernon et al., 2003). The outcomes of frequency specificity amongst them are inconsistent, showing both frequency-specific (Liu et al., 2022; Vernon et al., 2003) and non-specific (Dessy et al., 2020) effects. However, for non-relative SMR or Beta NFB training, some evidence points towards non-specific effects in the frequency domain (Chen & Lin, 2020; Pimenta et al., 2018), in line with our findings.

Interestingly, the spatial specificity of SMR NFB training differed between the two clusters of learners. Learners in Cluster 1 decreased SMR bilaterally across the targeted electrodes and equivalent electrodes in the opposite hemisphere. The decrease lasted until the fourth block and was then maintained with a slight lateralization towards the target hemisphere during the last block of NFB training. Cluster 2 learners displayed a spatially specific effect that lasted throughout the session and was accompanied by a continuous decrease. The non-learner showed a decrease focused on the target electrodes during the last two blocks, suggesting that learning took place, albeit not sufficient for being labelled a learner. Both learner clusters effectively down-regulated SMR; however, the selective and persistent down-regulation of SMR for Cluster 2, both in spatial and frequency domains, suggests that this group learned a more effective strategy.

Overall, Beta down-regulation was not as spatially selective to the targeted central electrodes as SMR, but seemed instead to be driven by power reductions in peripheral areas ranging from frontal to posterior regions. Most participants (75%), classified as Cluster 2 learners, displayed spatial dynamics resembling those of SMR Cluster 1, with strong bilateral decreases across sensorimotor regions. Also considering the lack of frequency specificity for Beta regulation, we suggest that central Beta was not directly modulated through the NFB, but indirectly through other features, perhaps more alike the SMR features used in this study. In contrast to occipital Alpha, our results show that central Beta NFB would require a stricter protocol in terms of selective neural self-regulation in frequency and spatial domains.

### Frequency and spatial specificity of cortical self-regulation

In this study, we employed a non-selective approach to NFB regulation, meaning that we did not penalize irrelevant activity modulations. While this is a widely used approach (Gruzelier, 2014; Rogala et al., 2016), the specificity of the neural self-regulation is rarely studied (Zoefel et al., 2011). Furthermore, imposing excessive constraints on the feedback signal (e.g., multiple inhibitory bands) may reduce the contingency of the feedback and hinder operant learning (Strehl, 2014). It is interesting to note that a portion of the learners were able to selectively enhance occipital alpha in both the frequency and spatial domains, whereas none of the learners could selectively reduce central Beta or unilateral centrotemporal SMR in both frequency and spatial domains. Only a portion of SMR learners reduced power over unilateral centrotemporal areas, whereas central Beta learners were non-selective in neither the spatial nor the frequency domain. Alpha up-regulation using an NFB protocol that does not constrain modulations in other frequencies has been shown to still produce frequency-selective outcomes (Grosselin et al., 2021), in line with what our results are showing. At the same time, non-selective protocols in Beta and SMR enhancing NFB have been shown to produce non-specific power modulations in a wide range of frequencies (Ghoshuni et al., 2012; Kober et al., 2017). There is limited knowledge in terms of the specificity in reducing SMR or Beta through NFB, but our results point to similar non-specific modulations, in line with one preliminary report (Dussard et al., 2024). It is possible that SMR and Beta power are associated with different mental strategies, some of which are related to more broadband reductions across large cortical areas (Egner et al., 2004). Therefore, effectively targeting a specific behavior or cognitive process through SMR or Beta NFB training may require protocols that are selective in both frequency and spatial domains.

### Limitations and Future Directions

The cluster-specific findings, particularly for non-learners, should be interpreted with caution due to limited sample sizes. These results are best viewed as hypothesis-generating observations rather than evidence of stable subgroup structure. Nevertheless, the relative frequency of the observed learning dynamics may help inform power calculations and recruitment targets for future studies investigating subgroup-specific neurofeedback trajectories.

A critical limitation of the present neurofeedback design is the lack of explicit constraints on adjacent frequency bands. The observed reductions in SMR and Beta power were consistently accompanied by decreases across other frequency bands, indicating that the obtained effects were not frequency-specific. This pattern suggests that participants may have modulated a more global cortical state (e.g., arousal or attentional engagement) rather than achieving spectrally specific control over the targeted rhythms. Importantly, this limitation arises from the non-selective design of the protocol, which did not penalize off-target activity or incorporate relative power measures. Future protocols could incorporate relative power metrics or explicit penalization of adjacent frequencies to better disentangle frequency-specific regulation from broadband effects. These findings highlight a fundamental ambiguity in neurofeedback interpretation: successful task performance does not necessarily reflect targeted neural modulation, but may instead arise from non-specific changes in overall brain state.

Another limitation of the present study is that frequency band and regulation direction were not independently manipulated. Specifically, lower-frequency rhythms (Theta, Alpha) were trained using up-regulation, whereas higher-frequency rhythms (SMR, Beta) were trained using down-regulation. As a result, it is not possible to fully disentangle whether observed differences in performance reflect frequency-specific properties or more general asymmetries between up- and down-regulation. This design reflects a deliberate trade-off. The primary aim of the study was to enable a controlled within-subject comparison across multiple commonly used neurofeedback features. Orthogonally manipulating both frequency and regulation direction would have substantially increased the number of experimental conditions, reducing statistical power within each condition given the available sample size. We therefore prioritized breadth of feature comparison while maintaining sufficient observations per condition. Importantly, this limitation does not affect the central conclusion that neurofeedback performance varies within individuals across features. However, future studies should explicitly dissociate frequency and regulation direction to determine the extent to which these factors independently contribute to learnability.

### Conclusion

In summary, this study examined inter-individual variability in EEG neurofeedback learning across four frequency bands, employing both novel analytic methods and detailed characterization of self-reported strategies. Our results demonstrate that all participants were able to regulate at least two EEG features, and many of the participants were able to regulate three features. With these results, we challenge the concept of the universal “non-learner” and reframe non-performance as a feature-specific, rather than person-specific, phenomenon. The ability to up-regulate frontal midline Theta was uniformly absent, indicating that not all oscillatory targets are equally accessible to volitional control with our protocol. By contrast, Alpha, SMR, and Beta were successfully modulated by most participants, albeit with distinct cognitive strategies and varying degrees of spatial and frequency specificities.

Methodologically, this work advances neurofeedback research in four ways. First, we introduce a method for modeling non-linear dynamics in neurofeedback data. Second, we introduced a clustering approach to classify the shape of individual learning trajectories, enabling the identification of distinct regulation profiles rather than relying on aggregated group means or regression slopes. Third, we visualized and compared entire learning trajectories for these clusters, providing a more nuanced view of within-session dynamics. Fourth, we reported modulation across all major frequency bands, rather than restricting analyses to the trained band. This revealed cross-band interactions that are often not reported in the field.

These findings have significant implications for both basic science and clinical applications, advocating for a shift away from one-size-fits-all protocols towards a personalized approach. By identifying an individual’s neurofeedback aptitude profile, we can better tailor interventions to harness their specific capacity for neural self-regulation, potentially enhancing the efficacy and reach of this promising technology.

## Acknowledgment

This work was supported by a grant from The Knowledge Foundation to Elaine Astrand (KKS HÖG project #20190099).

## Data Availability Statement

The preprocessed data and analysis code required to reproduce the statistical results and figures of this study are available in the Zenodo repository via the following anonymous XXX. The contents of the repository should be kept confidential until the manuscript is published and the data are released to the general public. Specific inquiries regarding the raw data may be directed to the corresponding author.

## Ethics approval

The study adhered to the principles of the Declaration of Helsinki and complied with local rules and regulations, as approved by the Swedish Ethical Review Authority (Dnr. 2021-03121).

